# Denoising Genome-wide Histone ChIP-seq with Convolutional Neural Networks

**DOI:** 10.1101/052118

**Authors:** Pang Wei Koh, Emma Pierson, Anshul Kundaje

## Abstract

**Motivation:** Chromatin immunoprecipitation sequencing (ChIP-seq) experiments are commonly used to obtain genome-wide profiles of histone modifications associated with different types of functional genomic elements. However, the quality of histone ChIP-seq data is affected by a myriad of experimental parameters such as the amount of input DNA, antibody specificity, ChIP enrichment, and sequencing depth. Making accurate inferences from chromatin profiling experiments that involve diverse experimental parameters is challenging.

**Results:** We introduce a convolutional denoising algorithm, Coda, that uses convolutional neural networks to learn a mapping from suboptimal to high-quality histone ChIP-seq data. This overcomes various sources of noise and variability, substantially enhancing and recovering signal when applied to low-quality chromatin profiling datasets across individuals, cell types, and species. Our method has the potential to improve data quality at reduced costs. More broadly, this approach – using a high-dimensional discriminative model to encode a generative noise process – is generally applicable to other biological domains where it is easy to generate noisy data but difficult to analytically characterize the noise or underlying data distribution.

**Availability:** https://github.com/kundajelab/coda

## 1 Introduction

Distinct combinations of histone modifications are associated with different classes of functional genomic elements such as promoters, enhancers, and genes (Consortium *et al*., 2015). Chromatin immunoprecipitation followed by sequencing (ChIP-seq) experiments targeting these histone modifications have been used to profile genome-wide chromatin state in diverse populations of cell types and tissues (Consortium *et al*., 2015), allowing us to better understand the mechanisms of development (Bernstein *et al*., 2006) and disease (Gjoneska *et al*., 2015).

However, the quality of histone ChIP-seq experiments is affected by a number of experimental parameters including antibody specificity and efficiency, library complexity, and sequencing depth(Jung *et al*., 2014). Achieving optimal experimental parameters and comparable data quality across experiments is often difficult, costly, or even impossible, resulting in low sensitivity and specificity of measurements especially in low input samples such as rare populations of primary cells and tissues (Brind’Amour *et al*., 2015; Cao *et al*., 2015; Acevedo *et al*., 2007). For example, (Brind’Amour *et al*., 2015) found that single mouse embryos do not provide enough cells to profile using conventional ChIP-seq techniques. Similarly, (Acevedo *et al*., 2007) notes that tumor biopsies, fractionated mixed cell populations, and differentiating embryonic stem cells provide very small numbers of cells to use as input populations. Further, the high sequencing depths (>50-100M reads) required for saturated detection of enriched regions in mammalian genomes for several broad histone marks (Jung *et al*., 2014) are often not met due to cost and material constraints. Suboptimal and variable data quality significantly complicate and confound integrative analyses across large collections of data.

To overcome these limitations, we introduce here a convolutional denoising algorithm, called Coda, that uses convolutional neural networks (CNNs) (Jain and Seung, 2009; Krizhevsky *et al*., 2012) to learn a generalizable mapping between ‘suboptimal’ and high-quality ChIP-seq data (Fig. 1). Coda substantially attenuates three primary sources of noise– due to low sequencing depth, low cell input, and low ChIP enrichment– enhancing signal in low-quality samples across individuals, cell types, and species. Our approach is conceptually related to the existing literature on structured signal recovery, in particular supervised denoising in images (Jain and Seung, 2009; Xie *et al*., 2012; Mousavi *et al*., 2015) and speech (Maas and Le, 2012). It complements other efforts to impute missing genomic data, such as ChromImpute (Ernst and Kellis, 2015), which predict profiles for a missing target mark in a target cell type (e.g., H3K4me3 in embryonic stem cells) by leveraging other available marks in the target cell type (e.g., H3K27ac in embryonic stem cells) and target mark datasets in other reference cell types (e.g., H3K4me3 in 100s of other celltypes). In contrast, our models take in low-quality signal of multiple target marks in a target cell type and denoise them all (e.g., using low-quality H3K27ac and H3K4me3 signal from a given cell population to produce higher-quality H3K27ac and H3K4me3 signal in that same cell population).

**Fig. 1.**
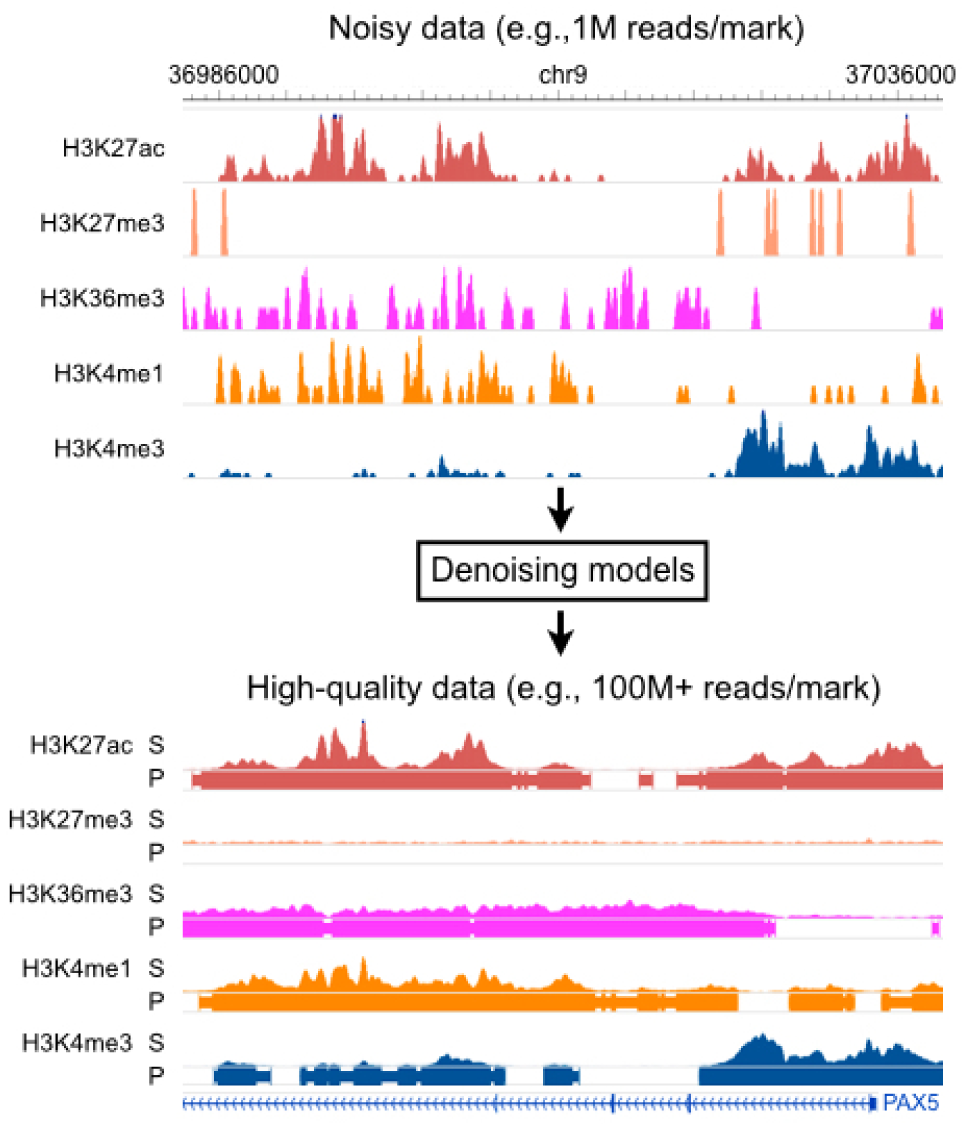
Overall model. Coda learns a transformation from noisy histone ChIP-seq data toa set of clean signal tracks and accurate peak calls. Top: a noisy signal track derived from 1M ChIP-seq reads per histone mark on the lymphoblastoid cell line GM12878. Bottom: a high-quality signal track derived from 100+M ChIP-seq reads per histone mark from the same experiment. S = Signal, P = Peak calls.

Neural networks have been successfully used to reduce noise in image data (Jain and Seung, 2009) and speech data (Maas and Le, 2012; Amodei *et al*., 2016), and there are several reasons to believe that neural networks could similarly denoise histone ChIP-seq data. First, histone marks have regular structure: peaks in each mark, for example, might tend to have certain widths and certain shapes. This means that a noisy signal can be denoised by a model that encodes prior expectations of what a clean signal should look like, just as humans use the regular structure in speech to decode noisy speech signals. Second, histone marks are correlated; thus, one noisy mark can be denoised using information from other noisy marks. Third, neural networks excel at flexibly learning complex nonlinear relationships when given large amounts of data, making them ideal for genome-wide applications. Indeed, neural networks have recently been successfully applied to many biological domains (Angermueller *et al*., 2016b): for example, they have been used to predict regulatory sequence determinants of DNA and RNA binding proteins (Alipanahi *et al*., 2015; Zhou and Troyanskaya, 2015), chromatin accessibility (Kelley *et al*., 2015), and methylation status (Angermueller *et al*., 2016a).

## 2 Methods

### Model

Coda takes in a pair of matching ChIP-seq datasets of the same histone modifications in the same cell-type – one high-quality and the other noisy– and uses convolutional neural networks (CNNs) to learn a mapping from the noisy to the high-quality ChIP-seq data. The noisy dataset used in training can be derived computationally (e.g., by subsampling the high-quality data) or experimentally (e.g., by conducting the same ChIP-seq experiment with fewer input cells). Once this mapping has been learned, the same mapping can then be applied to new, noisy data in any other cellular context with the same underlying noise structure.

For each type of noise (e.g., due to low cell numbers, sequencing depth, or enrichment) and each target histone mark, we train two separate CNNs to accomplish two tasks: a regression task to predict histone ChIP signal (i.e., the fold enrichment of ChIP reads over input DNA control) and a binary classification task to predict the presence or absence of a significant histone mark peak (Fig. 2). In total, if a given experiment has *M* marks, then we train 2*M* models separately (one regression and one classification model for each mark). Each individual model optionally makes use of the noisy ChIP-seq data from all available marks but outputs only one target histone mark. This allows us to learn separate features for each mark and task while still leveraging information from multiple input histone marks; we find empirically that this improves performance.

**Fig. 2.**
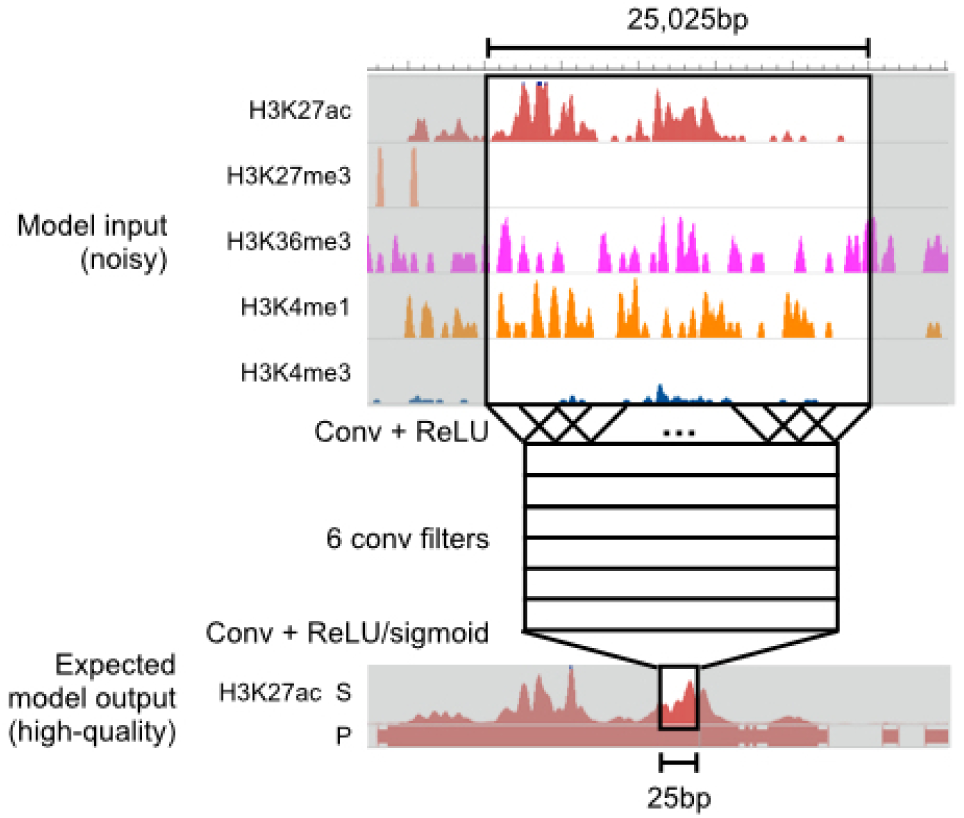
Model architecture. Coda learns two separate convolutional neural networks(CNN) for each target histone mark, one for regression (signal track reconstruction) and the other for classification (peak calling). All networks share the same architecture. Here, we show a schematic of a model trained to output a denoised signal track for H3K27ac. To make a prediction on a single location, we take in 25,025bp of data from all available histone marks centered at that location and pass it through two convolutional layers.

For computational efficiency, we first bin the genome into 25bp bins, averaging the signal in each bin. Let *L* be the number of bins in the genome (i.e., the length of the genome divided by 25). Each individual model takes in an *M* × *L* input matrix **X** and returns a 1 × *L* output vector **Y** representing the predicted high-quality signal (in the regression setting) or peak calls (in the classification setting). It does this by feeding the noisy data through a first convolutional layer, a rectified linear unit (ReLU) layer, a second convolutional layer, and then a final ReLU or sigmoid layer (for regression or classification, respectively). For the first convolutional layer, we use 6 convolutional filters, each 51 bins in length; for the second convolutional layer, we use a single filter of length 1001. Effectively, this means that a prediction at the *i*-th bin is a function of the noisy data at a 25,025bp window centered on the *i*-th bin.

The convolutional nature of our models (and the lack of max-pooling layers commonly seen in neural network architectures for computer vision) enables us to do efficient genome-wide prediction, as 98% of the computation required for predicting signal at the *i*-th bin is shared with the computation required for predicting the (*i* + 1)-th bin. In particular, to compute the prediction at the i-th bin, the network needs to perform 6 × 1001 × 51 operations at the first convolutional layer and 6 × 1001 operations at the second convolutional layer. To compute the prediction at the (*i*+1)-th bin, the network needs to perform only 6 × 51 more operations at the first convolutional layer and 6 × 1001 operations at the second convolutional layer, saving 6 × 1001 × 50 operations. Other models, especially non-linear models such as random forests, would require a completely separate set of computations for each bin and are therefore significantly more computationally expensive when it comes to making predictions across the entire genome.

### Training and evaluation

We applied Coda to three distinct sources of noise: low sequencing depth, low cell input, and low ChIP enrichment. In all cases, the inputs to our model were noisy signal measurements of multiple histone marks (see Data Availability and Processing for more details), and we trained separate models to predict the high-quality signal and peak calls for each target mark.

For the regression tasks (predicting signal), we evaluated performance by computing the Pearson correlation and mean squared error (MSE) between the predicted and measured high-quality fold-enrichment signal profiles after an inverse hyperbolic sine transformation, which reduced the dominance of outliers. We compared this to the baseline performance obtained by directly comparing the noisy and high-quality signal profiles of the target mark (after the same inverse hyperbolic sine transformation).

For the classification tasks (predicting presence or absence of a peak), we compared our model’s output to peaks called by the MACS2 peak caller (Feng *et al*., 2012) on the high-quality signal for the target mark. As our dataset is unbalanced – peaks only make up a small proportion of the genome – we evaluated performance by computing the area under the precision-recall curve (AUPRC), a standard measure of classification performance for unbalanced datasets (Davis and Goadrich, 2006). We compared the AUPRC of our model to a baseline obtained by comparing MACS2 peaks on the noisy data for the target mark to those obtained from the high-quality data for the target mark (see Data Availability and Processing for further details on dataset preparation).

We trained our models on 50,000 positions randomly sampled from peak regions of the genome and 50,000 positions sampled from non-peak regions, sampling from each autosome with equal likelihood. We defined peak regions using the output mark of interest and with the high-quality data. Further increasing dataset size did not increase performance; as each sample covered 25,025bp, 100,000 samples had good coverage of the entire genome. We selected the training dataset to be balanced because a uniformly drawn dataset would have had very few peaks, making it difficult for the model to learn to predict at peak regions; however, the test results reported in this paper are on the entire (unbalanced) genome. We used the *Keras* package (François Chollet, 2015) for training and AdaGrad (Duchi *et al*., 2011) as the optimizer, stopping training if validation loss did not improve for three consecutive epochs. We did not observe overfitting with our models (train and test error were comparable), and therefore opted not to use common regularization techniques such as dropout (Srivastava *et al*., 2014).

We chose model hyperparameters and architecture through hold-out validation on the low-sequencing-depth denoising task with GM12878 as the training cell line (Kasowski *et al*., 2013), holding out a random 20% subset of the training data for validation; this task will be discussed in more detail in the next section. The model architecture described above (6 convolutional filters each 51 bins in length in the first layer, and 1 convolutional filter of length 1001 in the second layer) yielded optimal validation performance out of the configurations we tried (varying the number of convolutional filters and the lengths of the filters by up to an order of magnitude). Adding an additional layer to the neural network brought a modest increase in performance at the cost of more computation time and complexity. To be sure that our model architecture generalized, we used the same architecture and hyperparameters for all denoising tasks without any further tuning.

## 3 Results

### Removing noise from low sequencing depth data

A minimum of 40-50M reads is recommended for optimal sensitivity for histone ChIP-seq experiments in human samples targeting most canonical histone marks (Jung *et al*., 2014). As adhering to this standard can often be infeasible due to cost and other limitations, a substantial proportion of publicly available datasets do not meet these standards. Motivated by these constraints, we tested whether our model could recover high-read depth signal from low-read depth experiments.

#### Training and testing on the same cell type across different individuals

We evaluated Coda on lymphoblastoid cell lines (LCLs) derived from six individuals of diverse ancestry (European (CEU), Yoruba (YRI), Japanese, Han Chinese, San) (Kasowski *et al*., 2013). We used the CEU-derived cell line (GM12878) to train our model to reconstruct the high-depth signal (100M+ reads per mark; exact numbers in Data Availability and Processing) from a simulated noisy signal derived by subsampling 1M reads per mark. On the other five cell lines, Coda significantly improved Pearson correlation between the full and noisy signal (Fig. 3A, left) and the accuracy of peak calling (Fig. 3A, right). Using just 1M reads per mark, the predicted output of our model was equivalent in quality to signal derived from 15M+ reads (H3K27ac) and 25M+ reads (H3K36me3) (Fig. 3B). Fig. 4 shows how Coda can accurately reconstruct histone modification levels at the promoter of the *PAX5* gene, a master transcription factor required for differentiation into the B-lymphoid lineage (Nutt *et al*., 1999).

**Fig. 3.**
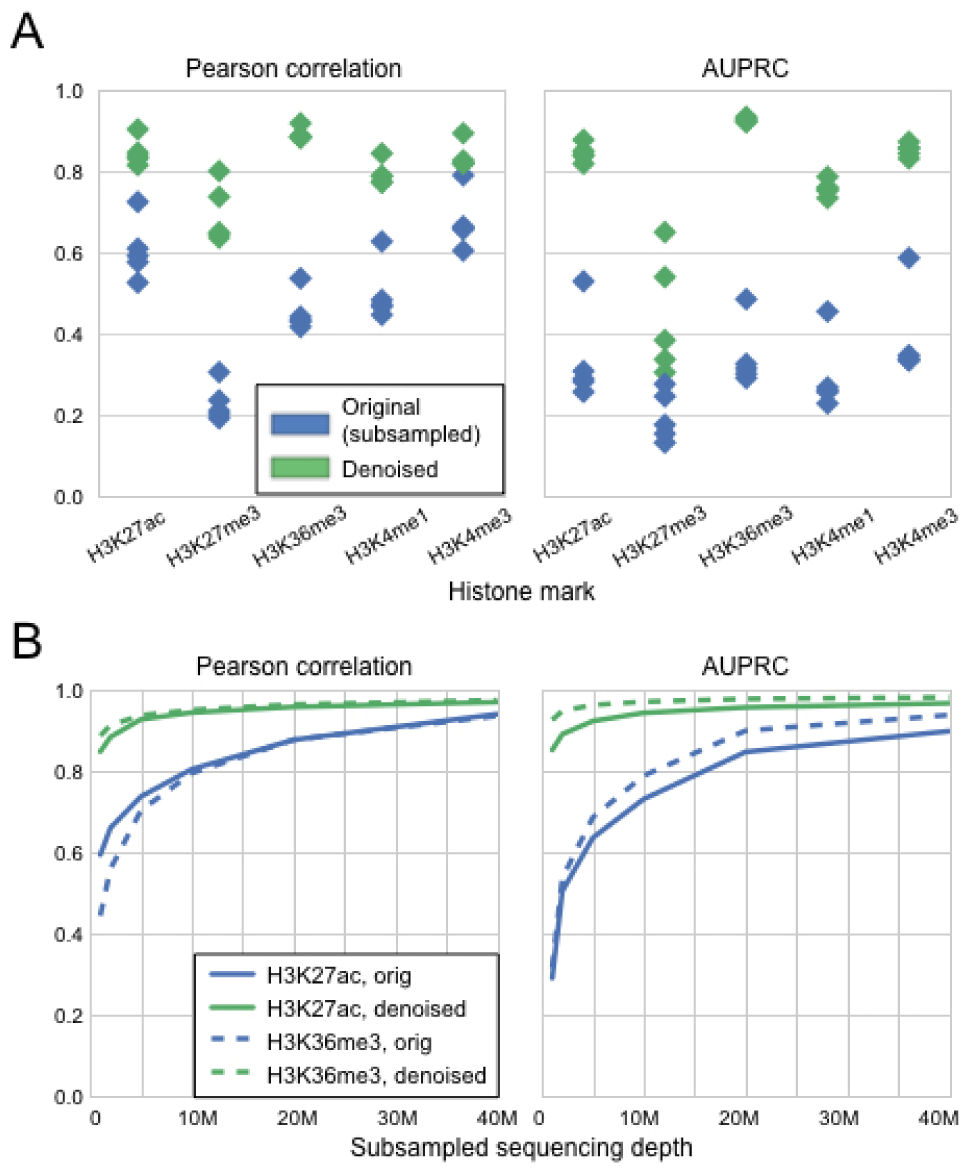
Coda removes noise from low-sequencing-depth experiments onlymphoblastoid cell lines derived from different individuals. (A) Compared to the signal from subsampled reads (blue), the denoised signal (green) shows greater correlation with the full signal (left) and more accurate peak-calling (right) across all cell lines. The model was trained on GM12878 and tested on different cell lines; within each column in the plot, each point is a single test cell line. (B) With 1M reads per mark, the denoised H3K27ac data is equivalent in quality to a dataset with 15M+ reads per mark, and the H3K36me3 data is equivalent in quality to a dataset with 25M+ reads per mark. Similar results hold for other marks. These results are from training on GM12878 and testing on GM18526.

**Fig. 4.**
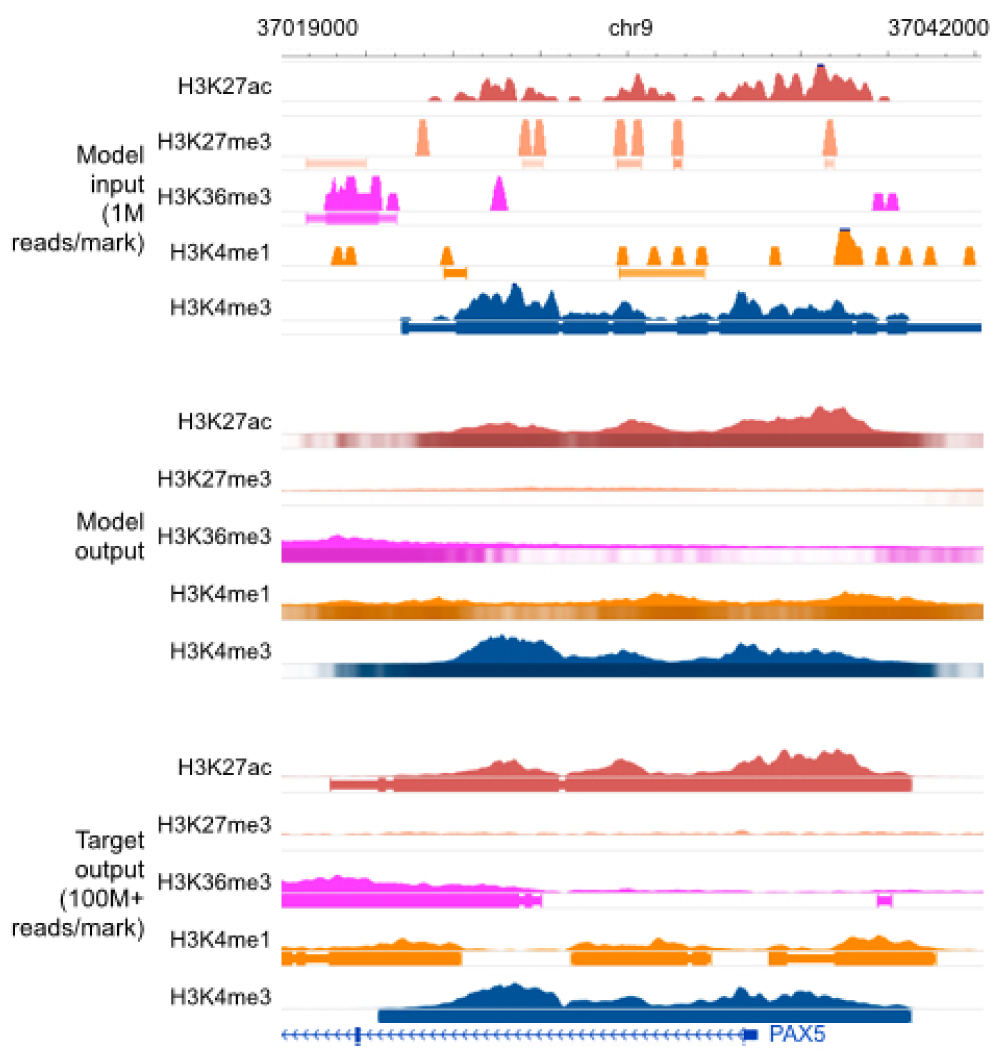
Genome browser tracks for low-sequencing-depth experiments. We comparenoisy signal and peak calls obtained from 1M reads per mark (top) with Coda’s output (middle) and the target, high-quality signal and peak calls obtained from 100M+ reads per mark (bottom) at the PAX5 promoter. Coda successfully cleans up signal across all histone marks and correctly calls the H3K27ac, H3K36me3, and H3K4me1 peaks (missed in the noisy data) while removing the spurious H3K27me3 peak calls. Note that we show the noisy peak calls to allow for comparisons; Coda uses only the noisy signal, not the peak calls, as input. The signal tracks are in arcsinh units, with the following y-axis scales: H3K27ac: 0-160, H3K27me3: 0-20, H3K36me3 and H3K4me1: 0-40, H3K4me3: 300. The shading of the peak tracks that the model outputs represent the strength of the peak call on a scale of 0-1.

We confirmed Coda was not simply memorizing the profile of the training cell line (GM12878) and copying it to the test cell lines by examining differential regions, called by DESeq (Anders and Huber, 2010), between GM12878 and the other cell lines (Kasowski *et al*., 2013). Coda improved correlation and peak-calling even in those regions (Table 1). Similarly, it also improved correlation on the regions of the genome with enriched signal, i.e., called as statistically significant peaks (Table 2).

**Table 1.** Denoising differential regions (diff. reg.) between test cell line GM18526 and training cell line GM12878. Performance reported is improvement of the denoised model over baseline (original, subsampled reads) on the test cell line.In parentheses we report the baseline results followed by the denoised results.Peak-calling results on H3K27me3 are omitted due to the lack of peak calls in differential regions; all results on H3K36me3 are omitted due to low number of differential regions.

|  | MSE (diff. reg.) | Pearson R (diff. reg.) | AUPRC (diff. reg.) |
| --- | --- | --- | --- |
| H3K4me1 | **-85%** (4.01, 0.57) | **+59%** (0.37, 0.59) | **+03%** (0.93, 0.97) |
| H3K4me3 | **-75%** (2.88, 0.70) | **+14%** (0.63, 0.72) | **+11%** (0.78, 0.87) |
| H3K27ac | **-86%** (3.43, 0.48) | **+39%** (0.55, 0.77) | **+06%** (0.90, 0.96) |
| H3K27me3 | **-80%** (0.78, 0.15) | **+106%** (0.14, 0.30) | - |

**Table 2.**
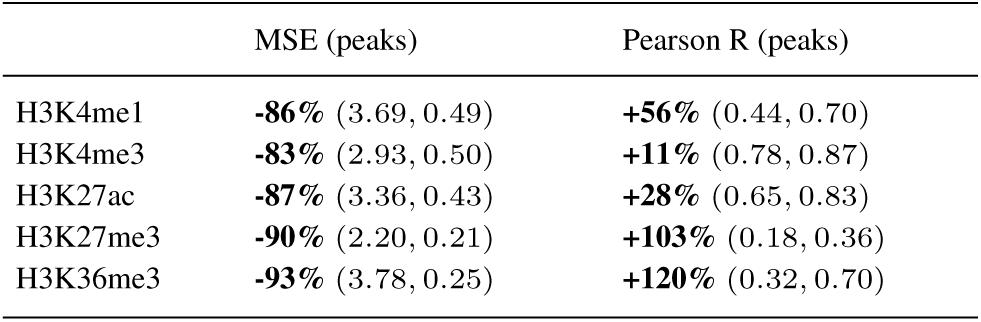
Denoising peak regions between test cell line GM18526 and training cell line GM12878. Performance reported is improvement of the denoised model over baseline (original, subsampled reads) on the test cell line. In parentheses we report the baseline results followed by the denoised results.

|  | MSE (peaks) | Pearson R (peaks) |
| --- | --- | --- |
| H3K4me1 | **-86%** (3.69, 0.49) | **+56%** (0.44, 0.70) |
| H3K4me3 | **-83%** (2.93, 0.50) | **+11%** (0.78, 0.87) |
| H3K27ac | **-87%** (3.36, 0.43) | **+28%** (0.65, 0.83) |
| H3K27me3 | **-90%** (2.20, 0.21) | **+103%** (0.18, 0.36) |
| H3K36me3 | **-93%** (3.78, 0.25) | **+120%** (0.32, 0.70) |

#### Training and testing on different cell types across different individuals

We next assessed if Coda could be trained on one cell type in one individual and used to denoise low-sequencing-depth data from a different cell type in a different individual. As above, the model was trained to output high-depth data (30M reads) from low-depth data (1M reads). We used histone ChIP-seq data spanning T-cells (E037), monocytes (E029), mesenchymal stem cells (MSCs, E026), and fibroblasts (E056) from the Roadmap Epigenomics Consortium (Consortium *et al*., 2015). Coda substantially improved the quality of the low-depth signal on the test cell type for all pairs of cell types (Table 3), illustrating its ability to denoise low-depth data on a cell type even if high-depth training data for that cell type is not available.

**Table 3.**
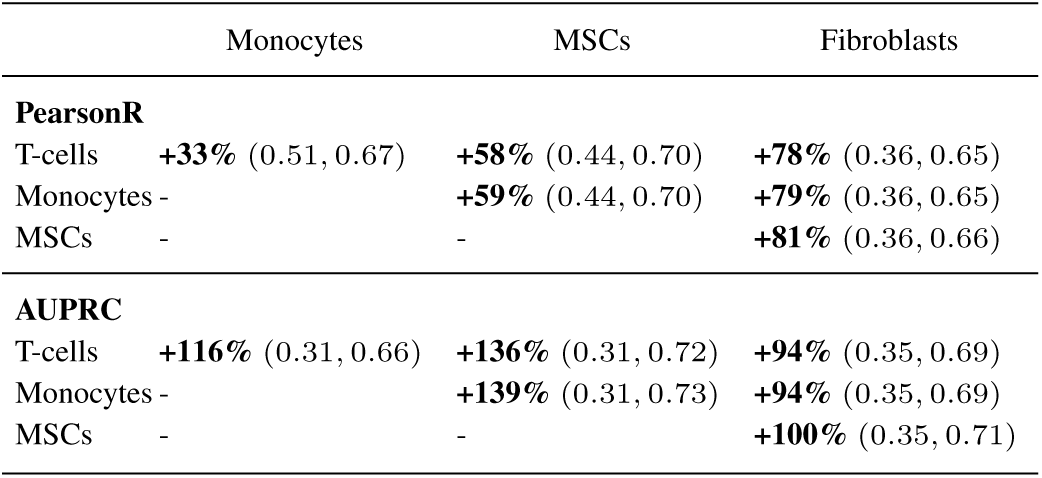
Cross cell-type experiments. Rows are train cell type, while columns are test cell type. In parentheses we report the baseline results followed by the denoised results, averaged across all histone marks used.

#### Coda outperforms linear baselines

We compared Coda to a linear and logistic regression baseline for signal denoising and peak calling, respectively. In both cases, we used an input region of the same size as Coda (i.e., 25,025bp centered on the location to be predicted, binned into 25bp bins). As noted above, the desire for computational efficiency in making genome-wide predictions across multiple marks limits the complexity of models that would be practically useful in genome-wide prediction.

When evaluated in the same cell type, different individual setting, Coda achieved 3x lower MSE on peak regions and 2x lower MSE on differential regions, with similar (very slightly better) MSE and correlation across the whole genome. This implies that Coda is better able to learn to match the exact values of the signal tracks on “difficult" regions (i.e., where there is the greatest deviation from the training signal), even though the linear model matches the rough shape. These regions are important to predict well because they can give insight into the differences between individuals and cell types.

We note that many forms of smoothing can be represented via linear regression. For example, a standard Gaussian filter can be interpreted as taking a linear combination of surrounding points with fixed coefficients. The comparison against a linear regression baseline therefore sets an upper bound for the performance of simple smoothing measures on this task (assuming no overfitting, which we do not observe in our case).

#### Comparisons to denoising and imputation

Next, we studied Coda’s performance in two additional settings: pure denoising (using the noisy target mark as the only input mark) and imputation from noise (using all noisy histone marks but the target mark as the input marks). This is in contrast to the standard setting described above, where we use all noisy histone marks, including the noisy version of the target mark, to recover a high-quality version of the target mark.

In the denoising case, Pearson correlation dropped by 0.03 points and AUPRC dropped by 0.05, on average, compared to when all marks were used as input. Thus, additional marks provided some information, but the denoised signal was still substantially better than the original subsampled signal.

In the imputation case, performance dropped somewhat on the narrow marks (H3K4me1, H3K4me3, H3K27ac; −0:12 correlation, −0:13 AUPRC) and dropped more on the broad marks (H3K27me3, H3K36me3; −0:29 correlation, −0:30 AUPRC). The gap in correlation was even larger within peak regions. Thus, having a noisy version of the target mark substantially boosts recovery of the high-quality signal.

### Removing noise from low cell input

Conventional ChIP-seq protocols require a large number of cells to reach the necessary sequencing depth and library complexity (Brind’Amour *et al*., 2015; Cao *et al*., 2015), precluding profiling when input material is limited. Several ChIP-seq protocols were recently developed to address this problem. We studied ULI-NChIP-seq (Brind’Amour *et al*., 2015) and MOWChIP-seq (Cao *et al*., 2015), which use low cell input (10^2^-10^3^ cells) to generate signal that is highly correlated, when averaged over bins of size 2-4kbp, with experiments with high cell input. However, at a finer scale of 25bp, the low-input signals from both protocols are poorly correlated with the high-input signals (Table 4).

**Table 4.** Low-cell-input experiments.We report improvement of the denoised model output over baseline (original low-input experiments), as compared to high-input experiments. In parentheses we report the baseline results followed by the denoised results.

|  | MSE | Pearson R | AUPRC |
| --- | --- | --- | --- |
| **ULI-NChIP** |  |  |  |
| H3K4me3 | **-61%** (1.39, 0.54) | **+208%** (0.13, 0.41) | **+61%** (0.24, 0.38) |
| H3K9me3 | **-46%** (0.51, 0.27) | **+28%** (0.41, 0.53) | **+32%** (0.28, 0.36) |
| H3K27me3 | **-41%** (0.68, 0.40) | **+57%** (0.34, 0.54) | **+32%** (0.34, 0.45) |
| **MOWChIP** |  |  |  |
| H3K4me3 | **-42%** (1.18, 0.68) | **+122%** (0.14, 0.31) | **+34%** (0.19, 0.25) |
| H3K27ac | **-21%** (1.44, 1.14) | **+159%** (0.09, 0.24) | **+66%** (0.15, 0.24) |

We thus used Coda to recover high-resolution, high-cell-input signal from low-cell-input signal specific to each protocol. For ULI-NChIP-seq, we used a single mouse embryonic stem cell dataset (Brind’Amour *et al*., 2015). For MOWChIP-seq, we trained on data from the human LCL GM12878 and tested on hematopoietic stem and progenitor cells (HSPCs) from mouse fetal liver (Cao *et al*., 2015). Coda successfully denoised the low-input signal from both protocols (Table 4). Fig. 5 illustrates our model denoising MOWChIP-seq signal across the *Runx1* gene, a key regulator of HSPCs (North *et al*., 2002); the results of peak calling were too noisy, even on the original 10,000-cell data, to allow for any qualitative judgment of improvement.

**Fig. 5.**
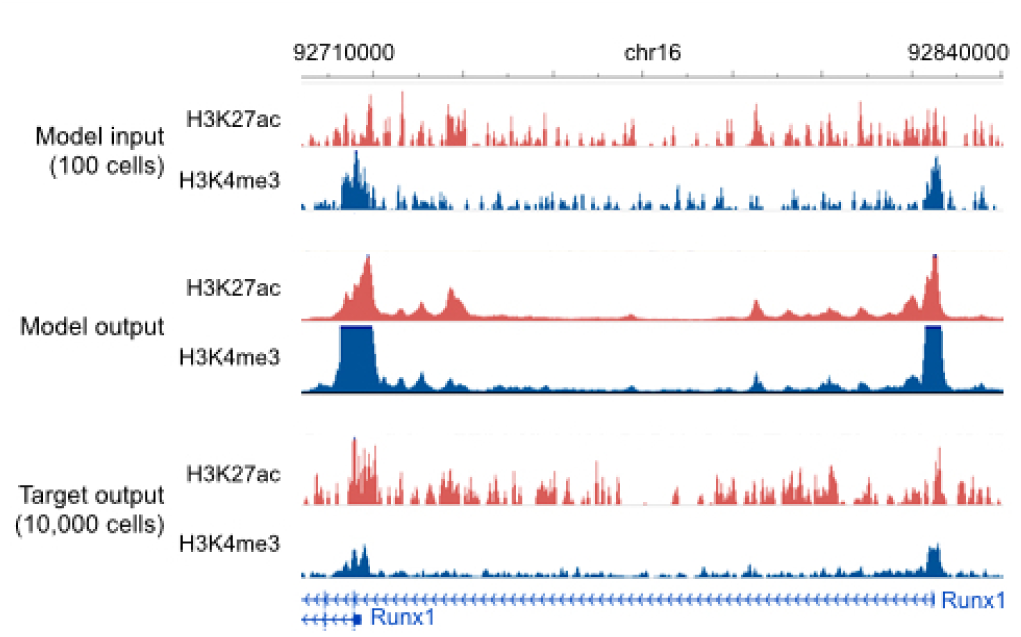
Genome browser tracks for low-cell-input experiments. We compare noisy signal obtained from 100 cells (top) with Coda’s output (middle) and the target, high-quality signal obtained from 10,000 cells (bottom) at the Runx1 gene in mouse hematopoietic stem and progenitor cells. The model was trained on MOWChIP-seq data generated from human LCL (GM12878) and captures two strong peaks at the promoters of the two isoform classes, removing much of the intervening noise. The signal tracks are in arcsinh units, with a scale of 0-40 for both histone marks.

We note that the Pearson correlations between the low cell input and high cell input in the original ULI-NChIP-seq (Brind’Amour *et al*., 2015) and MOWChIP-seq (Cao *et al*., 2015) papers are significantly higher than the ones we report here. We report lower correlations because we use a smaller bin size for the genome, as noted above; we look at correlation across the whole genome, instead of only at transcription start sites; and we compute correlation after an arcsinh transformation to prevent large peaks from dominating the correlation. Therefore, while the original low-cell-input data is suitable for studying histone ChIP-seq signal at a coarse-grained level and around genetic elements like transcription start sites, the denoised data is more accurate at a fine-grained level and across the whole genome.

### Removing noise from low-enrichment ChIP-seq

Histone ChIP-seq experiments use antibodies to enrich for genomic regions associated with the target histone mark. When an antibody with low specificity or sensitivity for the target is used, the resulting ChIP-seq data will be poorly enriched for the target mark. This is a major source of noise (Landt *et al*., 2012). We simulated results from low-enrichment experiments by corrupting GM12878 and GM18526 LCL data (Kasowski *et al*., 2013). For each histone mark profiled in those cell lines, we kept only 10% of the actual reads and replaced the other 90% with reads taken from the control ChIP-seq experiment, which was done without the use of any antibody; this simulates an antibody with very low specificity.

This corruption process significantly degraded the genome-wide Pearson correlation and the accuracy of peak calling (Table 5). This shows that recovering the true signal from the corrupted data cannot be achieved by simply linearly scaling the signal (e.g., multiplying the empirical fold enrichment by 10 since only 10% of the actual reads were kept), as if that were the case, the correlation would be unchanged. In contrast, when trained on GM12878 and tested on GM18526, Coda accurately recovered high-quality, uncorrupted signal from the corrupted data (Table 5). Fig. 6 shows a comparison of Coda’s output versus the corrupted and uncorrupted data at the promoter of the *EBF1* gene, another key transcription factor of the B-lymphoid lineage. (Nechanitzky *et al*., 2013)

**Table 5.** Low-enrichment experiments. We report improvement of the denoised model output over baseline (low-enrichment experiments), as compared to highenrichment experiments. In parentheses we report the baseline results followed by the denoised results.

|  | MSE | Pearson R | AUPRC |
| --- | --- | --- | --- |
| H3K4me1 | **-75%** (0.35, 0.09) | **+42%** (0.64, 0.91) | **+215%** (0.29, 0.92) |
| H3K4me3 | **-86%** (0.44, 0.06) | **+54%** (0.58, 0.91) | **+94%** (0.49, 0.95) |
| H3K27ac | **-70%** (0.37, 0.11) | **+37%** (0.65, 0.90) | **+121%** (0.43, 0.94) |
| H3K27me3 | **-61%** (0.27, 0.10) | **+88%** (0.42, 0.78) | **+242%** (0.14, 0.49) |
| H3K36me3 | **-82%** (0.36, 0.06) | **+47%** (0.65, 0.95) | **+168%** (0.36, 0.98) |

**Fig. 6.**
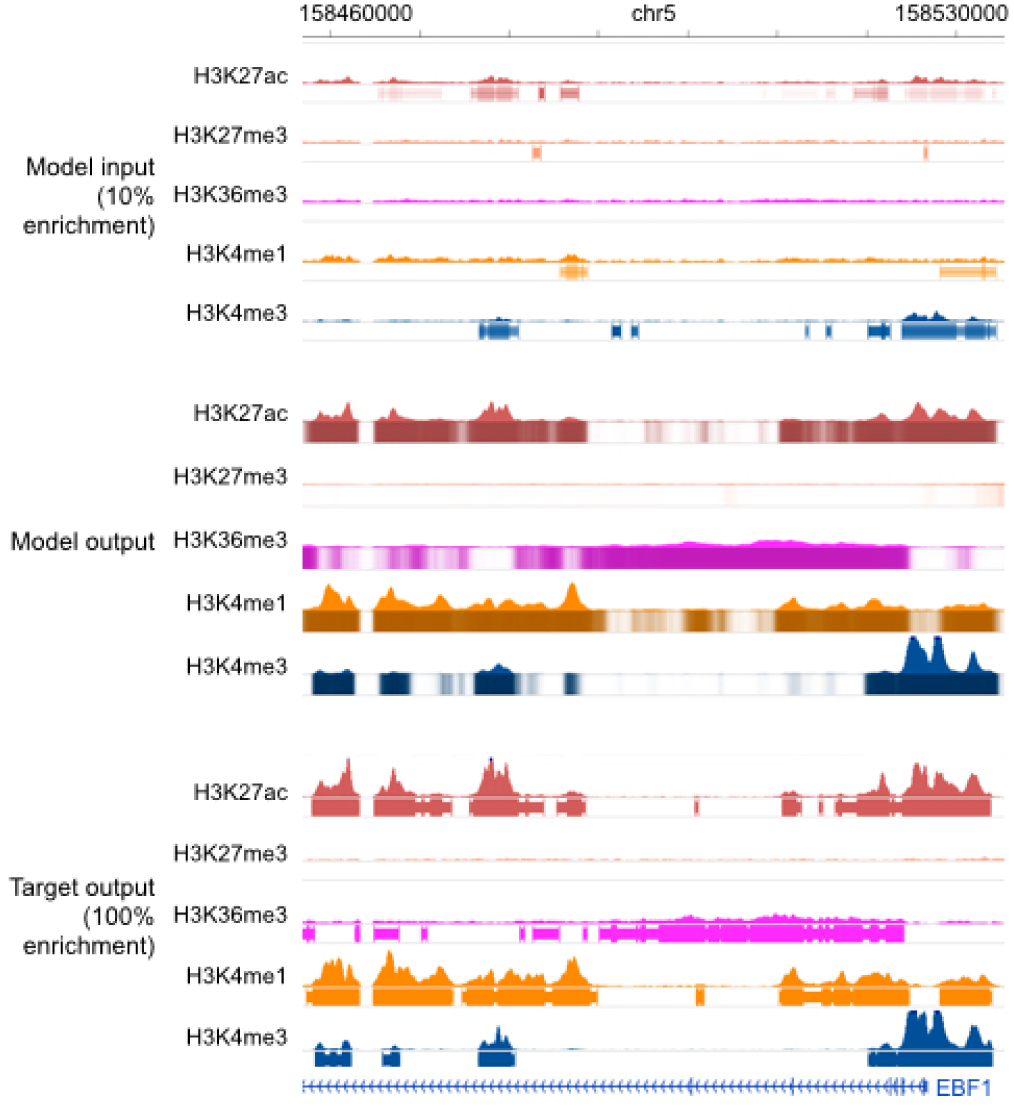
Genome browser tracks for low-enrichment ChIP-seq experiments. We compare noisy signal and peak calls obtained from the corrupted data with 10% enrichment (top) with Coda’s output (middle) and the target, high-quality signal and peak calls obtained from the uncorrupted data (bottom) at the EBF1 promoter. Coda significantly improves the signal-to-noise ratio and correctly calls the H3K27ac, H3K36me3, H3K4me1, and H3K4me3 peaks that were missed in the noisy data while removing a spurious H3K27me3 peak call. Note that we show the noisy peak calls to allow for comparisons; Coda uses only the noisy signal, not the peak calls, as input. The signal tracks are in arcsinh units, with the following y-axis scales: H3K27ac: 0-60, H3K27me3, H3K36me3, and H3K4me1: 0-40, H3K4me3: 100. The shading of the peak tracks that the model outputs represent the strength of the peak call on a scale of 0-1.

To further validate Coda’s output, we examined aggregate histone ChIP-seq signal around known biological regions of interest. In particular, we used the fact that H3K4me1 and H3K27ac, known enhancer marks, are enriched at DNase I hypersensitivity sites (DHSs), whereas H3K27me3 is depleted at DHSs. (Shu *et al*., 2011) For each of those marks, we compared the average uncorrupted signal, the average denoised signal, and the average low-enrichment signal within 5000 bp of the summits of DNase I hypersensitive peaks in GM12878 from ENCODE data (Bernstein *et al*., 2012). As expected, the corrupted, low-enrichment signal was biased by the reads from the control experiment and had significantly lower fold enrichment of H3K4me1 and H3K27ac at DHSs, compared to the uncorrupted signal. In contrast, the denoised signal was significantly more enriched at DHSs than the corrupted signal, more closely resembling the uncorrupted signal. Conversely, the corrupted signal had higher levels of H3K27me3 at DHSs, whereas the denoised signal had low levels of H3K27me3 throughout the DHS, similar to the uncorrupted signal though without a dip at the peak summit (Fig. 7).

**Fig. 7.**
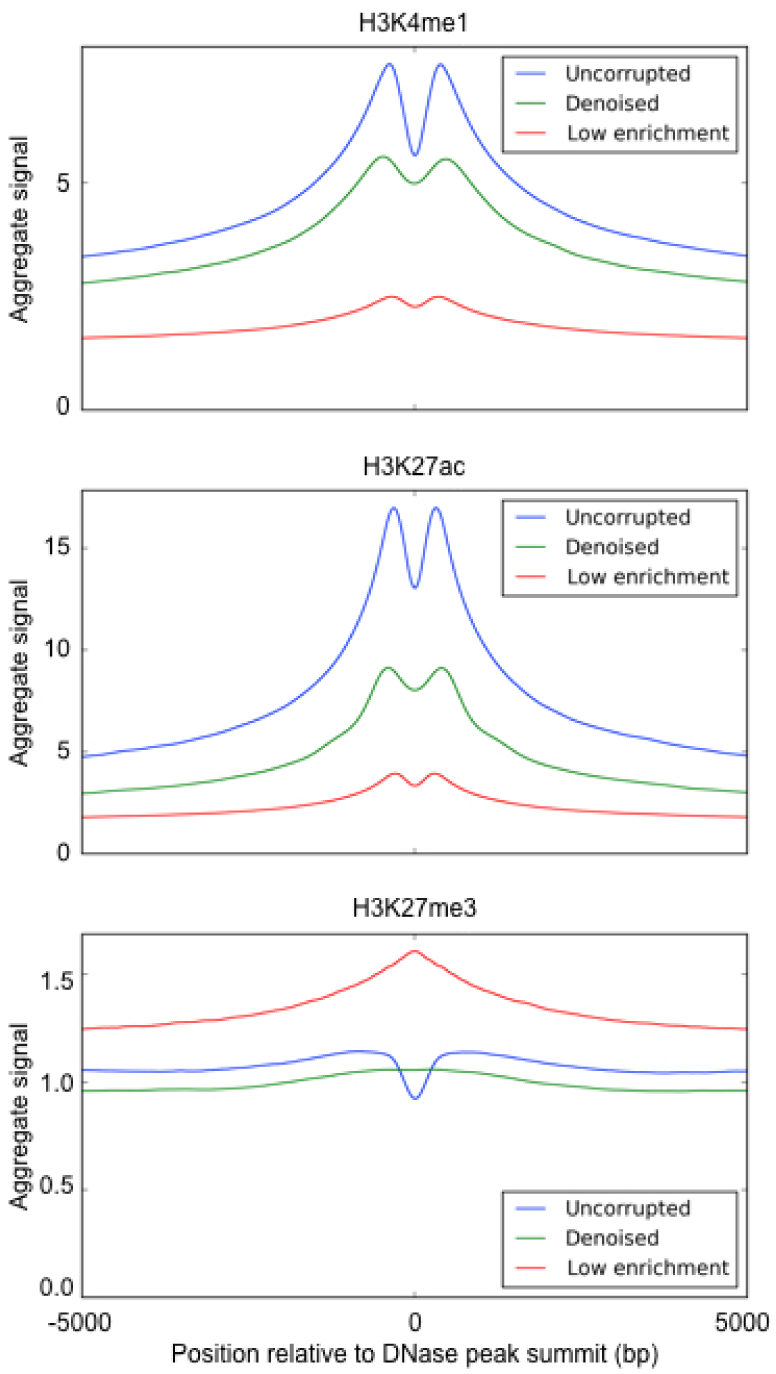
Aggregate histone ChIP-seq signal at DNase I hypersensitivity sites. We compare the average uncorrupted signal (Full), the average denoised signal (Denoised), and the average corrupted signal (Low enrichment) at DNase I hypersensitivity sites. Across all histone marks, the denoised signal is significantly more similar to the uncorrupted signal than the corrupted signal is.

## 4 Conclusion

We describe a convolutional denoising algorithm, Coda, that uses paired noisy and high-quality samples to substantially improve the quality of new, noisy ChIP-seq data. Our approach transfers information from generative noise processes (e.g., mixing in control reads to simulate low-enrichment, or performing low-input experiments) to a flexible discriminative model that can be used to denoise new data. We believe that a similar approach can be used in other biological assays, e.g., ATAC-seq and DNase-seq (Buenrostro *et al*., 2013; Crawford *et al*., 2006), where it is near impossible to analytically characterize all types of technical noise or the overall data distribution but possible to generate noisy versions of high-quality samples through experimental or computational perturbation. This can significantly reduce cost while maintaining or even improving quality, especially in high-throughput settings or when dealing with limited amounts of input material (e.g., in clinical studies).

An important caveat to our work is that the performance of Coda depends strongly on the similarity of the noise distributions and underlying data distributions in the test and training sets. For example, Coda expects that the relationships between different histone marks should be conserved between the test and training set. Thus, applying a set of trained Coda models to data that is very different from what it was trained on is unlikely to work. We also assume that the noise parameters in the test data are known in advance, e.g., the sequencing depth, the number of input cells, or the level of ChIP enrichment. In some cases (e.g., the low-sequencing-depth and low-cell-input settings) this is true, but in others (e.g., the low-enrichment setting) it is not always possible. An important direction for future work is to make Coda more robust; for example, training a single model over various settings of the noise parameters and various cell types could improve the generalizability of the models.

To further improve performance, more complex neural network architectures could also be explored: for example, using recurrent neural networks (Sutskever *et al*., 2014) to explicitly model long-range spatial correlations in the genome; multi-tasking across output marks instead of training separate models for each mark; or using deeper networks.

Another avenue for future work is exploring using more than just the noisy histone ChIP-seq data at test time. In this work, we use only the noisy data at test time, training our models to transform it into high-quality data. In reality, at test time we might have access to other data; for example, we might also have the DNA sequence of the test sample or access to high-quality ChIP-seq data on a closely related cell type. Other work has used DNA sequence to predict transcription factor binding (Alipanahi *et al*., 2015; Zhou and Troyanskaya, 2015), chromatin accessibility (Kelley *et al*., 2015), and methylation status (Angermueller *et al*., 2016a). A natural next step would be to combine the ideas from these methods with ours, e.g., by having a separate convolutional module in our neural network that incorporates sequence information and joins with the ChIP-seq module at an intermediate layer. Others have also used high-quality ChIP-seq data from closely related cell types for imputation (Ernst and Kellis, 2015); combining this with our denoising approach could help to avoid a potential pitfall of imputation approaches, namely the loss of cell-type-specific signal, while improving the accuracy of our denoised output.

Below, we provide a link to a script that trains a model for low-sequencing-depth noise using the LCL data described above. Since the type of noise can vary from context to context, we also provide the code for the general Coda framework to allow for developers of new protocols (e.g., new low-cell-count techniques) or core facilities that have high throughput to train Coda with data specific to their context.

### Data Availability and Processing

#### Datasets

We used the following publicly-available GEO datasets in this work:

1. GSE50893 for ChIP-seq data on LCLs (Kasowski *et al*., 2013)
2. GSE63523 for ULI-NChIP-seq data (Brind’Amour *et al*., 2015)
3. GSE65516 for MOWChIP-seq data (Cao *et al*., 2015)
4. GSM736620 for DNase I hypersensitive peaks (Bernstein *et al*., 2012)

For the low-sequencing-depth experiments, the full depth for GM12878 (training set) was 171M (million reads) for H3K4me1, 168M for H3K4me3, 328M for H3K27ac, 265M for H3K27me3, and 123M for H3K36me3. The full depth for GM18526 (test set) was 120M for H3K4me1, 136M for H3K4me3, 125M for H3K27ac, 138M for H327me3, and 223M for H3K36me3.

For the cross-cell-type experiments, we used the consolidated Roadmap Epigenomics data (Consortium *et al*., 2015), which is publicly available from http://egg2.wustl.edu/roadmap/data/byFileType/alignments/. Each mark is downsampled to a maximum of 30M reads to maximize consistency across marks; we used this as the full depth data, and downsampled to 1M reads for the noisy data. A detailed description of this dataset is available in (Roa, 2015).

#### Dataset preparation

##### Fold change signal profiles and peak calling

For each experiment, we used align2rawsignal (Kundaje, 2013) to generate signal tracks and MACS2 (Feng *et al*., 2012) to call peaks, as implemented in the *AQUAS* package (Lee and Kundaje, 2016). For the signal track, we used fold change relative to the expected uniform distribution of reads after an inverse hyperbolic sine transformation (Hoffman *et al*., 2012). We used the gappedPeaks output from MACS2 as the peak calls. For computational efficiency, we binned the genome into 25bp segments, averaging the signal in each segment.

We evaluated our peak calling on a bin-by-bin basis, i.e., our model output one number for each bin representing the probability that that bin was a true peak, and we treated each bin as a separate example for the purposes of computing AUPRC, our metric for peak calling performance. To get ground truth data for our peak calling tasks, we labeled each bin as “peak” or “non-peak” based on whether that bin was part of a peak called by MACS2 on the high-quality data.

Computing AUPRC requires predictions to be ranked in order of confidence. For our model, we used the output probabilities for each bin to calculate the ranking. MACS2 outputs both a peak p-value track, assigning a p-value to each genomic coordinate, and a set of binary peak calls. To measure baseline performance on the noisy data, we ranked each bin by the maximum peak p-value assigned by MACS2 to a genomic coordinate in that bin, unless that bin did not intersect with any of the binary peak calls, in which case it was assigned a p-value of – inf (i.e., ranked last). We did this to ensure that the high-quality peak track had an AUPRC of 1; empirically, this also improved performance of the noisy MACS2 baseline.

##### Histone marks used

We used different sets of input and output histone marks for different experiments depending on which marks each dataset provided. For the same cell type, different individual experiments (using lymphoblastoid cell lines), we trained and tested on H3K4me1, H3K4me3, H3K27ac, H3K27me3, and H3K36me3; we used the same data for the low-ChIP-enrichment experiments. For the different cell type, different individual experiments (using the uniformly-processed Roadmap Epigenomics Consortium datasets (Consortium *et al*., 2015)), we trained and tested on H3K4me1, H3K4me3, H3K9me3, H3K27ac, H3K27me3, and H3K36me3. For all of the above experiments, we also used data from the control experiments (no antibody) as input. Lastly, for the low-cell-input experiments, we used H3K4me3, H3K9me3, and H3K27me3 from the ULI-NChIP-seq dataset and H3K4me3 and H3K27ac from the MOWChIP-seq dataset.

##### Low-cell-input datasets

The ULI-NChIP-seq (Brind’Amour *et al*., 2015) and MOWChIP-seq (Cao *et al*., 2015) papers provided several datasets corresponding to different numbers of input cells used. For each protocol, we used the datasets with the lowest number of input cells as the noisy input data (ULI-NChIP-seq: 10^3^ cells for H3K9me3 and H3K27me3, 5*x*10^3^ cells for H3K4me3; MOWChIP-seq: 10^2^ cells) and the datasets with the highest number of input cells as the gold-standard, high-quality data (ULI-NChIP-seq: 10^6^ cells for H3K9me3, 10^5^ cells for H3K4me3 and H3K27me3; MOWChIP-seq: 10^4^ cells). The ULI-NChIP-seq data had matching low and high-input experiments only for a single cell type, so we divided it into chr5-19 for training, chr3-4 for validation, and chr1-2 for testing.

##### Code, data, and browser track availability

Our code is available on Github at https://github.com/kundajelab/coda, including a script that downloads pre-processed data and replicates the low-sequencing-depth experiments described above, as well as code for processing new data.

The figures of browser tracks (Figures 4, 5, and 6) shown above were taken from the Wash U Epigenome Browser (Zhou and Wang, 2012). Links to the entire browser tracks are as follows:

- Fig. 4, low-sequencing-depth experiments on LCL GM12878: http://epigenomegateway.wustl.edu/browser/?genome=hg19&session=KZvYzGBt03&statusId=107864126
- Fig. 5, low-cell-count experiments on mouse HSPCs: http://epigenomegateway.wustl.edu/browser/?genome=hg19&session=KZvYzGBt03&statusId=107864126
- Fig. 6, low-enrichment experiments on LCL GM12878: http://epigenomegateway.wustl.edu/browser/?genome=hg19&session=3hDZdGiGmF&statusId=1913128468

## Acknowledgements

We thank Jin-Wook Lee for his assistance with the AQUAS pipeline and Kyle Loh, Irene Kaplow, and Nasa Sinnott-Armstrong for their helpful feedback and suggestions.

